# Parsimonious scenario for the emergence of viroid-like replicons *de novo*

**DOI:** 10.1101/593640

**Authors:** Pablo Catalán, Santiago F. Elena, José A. Cuesta, Susanna Manrubia

## Abstract

Viroids are small, non-coding, circular RNA molecules that infect plants. Different hypotheses for their evolutionary origin have been put forward, such as an early emergence in a precellular RNA World or several *de novo* independent evolutionary origins in plants. Here we discuss the plausibility of *de novo* emergence of viroid-like replicons by giving theoretical support to the likelihood of different steps along a parsimonious evolutionary pathway. While Avsunviroidae-like structures are relatively easy to obtain through evolution of a population of random RNA sequences of fixed length, rod-like structures typical of Pospiviroidae are difficult to fix. Using different quantitative approaches, we evaluate the likelihood that RNA sequences fold into a rod-like structure and bear specific sequence motifs facilitating interactions with other molecules, *e.g.* RNA polymerases, RNases and ligases. By means of numerical simulations, we show that circular RNA replicons analogous to Pospiviroidae emerge if evolution is seeded with minimal circular RNAs that grow through the gradual addition of nucleotides. Further, these rod-like replicons often maintain their structure if independent functional modules are acquired that impose selective constraints. The evolutionary scenario we propose here is consistent with the structural and biochemical properties of viroids described to date.

## 1. Introduction

Since their discovery in 1971 [1], viroids have elicited both amazement and attention. Despite a small size of a few hundred nucleotides (nt) and a non-coding RNA sequence, these RNA molecules behave as competent and persistent replicators in plants, to our current knowledge their unique natural hosts. The origin of viroids has been one of the most controversial issues regarding these small pathogens. Most proposals in the literature have sought an origin either related to other extant small RNAs (see a review of hypotheses in [2]) or, in the light of viroid properties, suggest a possible old origin in a precellular RNA World [3,4].

The sequence and the structure of existing viroids must have evolved as a response to a variety of selection pressures and evolutionary constraints to guarantee their successful replication and propagation. Different structural domains of viroids, occasionally overlapping in sequence [5], are related to a variety of functions such as cell-to-cell and systemic movements, replication, circularization, or pathogenicity, and to ribozyme-like activities such as self-cleavage of multimeric intermediates of replication into individual genomes [2,5–7]. While compact secondary structures (especially the rod-like fold characteristic of Pospiviroidae) have been identified as a constraint for viroid evolution [8], their preservation seems essential to avoid degradation and inactivation [2] and to minimize the effect of deleterious mutations [9,10]. Still, their rod-like structures are less stiff than typical double-stranded RNA (dsRNA), a feature that may as well play a functional role, as it may facilitate recognition by RNA polymerases that transcribe dsRNA templates into single-stranded RNAs (ssRNA) [4,7].

Evidence for rapid evolution in viroids is mounting. As early as in 1996, Theodor Diener noted that “Results from site-directed mutagenesis experiments indicate that, upon exposure to selective pressures, viroids can evolve extremely rapidly, with another, fitter, component of the quasi-species often becoming dominant within days or weeks.” [11]. Viroids do behave as quasispecies [12], meaning that new mutations should be fixed through evolution and co-evolution with their molecular environment, a dynamical process that has been amply documented in viruses [13]. Though the small genomes of viroids allow faithful replication despite high mutations rates [14], the latter differ significantly in the two viroid families. The spontaneous mutation rates estimated for Avsunviroidae are between 1/400 for CChMVd [14] and 1/800 for ELVd [15], but that of PSTVd —and potentially other Pospiviroidae— is lower (in the range 1/7000 to 1/3800), being comparable to the mutation rate of RNA viruses [15]. In any case, such high mutation rates entail a population heterogeneity that has been confirmed in recent years. In a study with PLMVd where a peach tree was infected with a clonal inoculum, almost 4,000 different sequences were identified after just six months of in-plant evolution [16]. Only about 50% of the positions were fully conserved, and sequences differing in approximately 50 mutations from the parental sequence (which was not recovered from the evolved population) were identified [16]. Significant variations in the consensus sequence of GYSVd have been described in relation to the life-history of the grapevines it infects [17], while sequence variability increased in HLVd as a result of sudden environmental changes [18]. Moreover, the amount and nature of variability in CEVd strongly depended on the citrus host variety infected and could revert from configurations when hosts are reverted [19]. The mounting number of observations speaking for the high heterogeneity of viroid populations, their small genomes, and their rapid sequence change strongly suggests that we might be in a difficult position to solve questions on viroid’s origin if evidence has to rely solely on sequence similarity. Actually, it was chiefly the lack of sequence homology [11] that led to the dismissal of hypotheses suggesting that viroids might have descended from a variety of cellular RNAs [2].

In some scenarios dealing with an old precellular origin of viroids, they are portrayed as RNA molecules with properties very similar to those of extant viroids [4,20], which happen to fit into the chemical conditions of an early RNA World. However, and in a sense analogous to Spiegelman’s monster [21], viroids behave as minimal replicators. The *in vitro* evolution of the RNA genome of the Q*β* phage, with an approximate length of 4,500 nt, led to the eventual selection of a replicating RNA molecule (the monster) with just 218 nt [21]. This experiment demonstrated that, given the appropriate environment, complex genomes may reduce the set of functions they perform (or encode) to the minimum that guarantees their persistence. That scenario is akin to a top-down approach where viroids could have started as complex replicators in a (perhaps) simple molecular environment, bearing a larger array of functions that were subsequently lost to yield their present conformation [3, 22]. A complementary conceptual scenario may also apply: In a bottom-up approach (or *de novo* origin), viroid-like replicators might have come into being as serendipitous replicating sequences that subsequently acquired additional functions from a surrounding complex molecular environment. This is the scenario that we aim to explore here.

The main focus of this contribution is to quantitatively examine to which extent several of the known features of recognized viroids are easy to obtain through evolution. Many of their sequence and structural features can be obtained as a response to a number of selection pressures that are easy to cast in a simple fitness function: requiring an average folding energy (such that the folded structure is stable but able to be opened) may affect at once the G+C content and the fraction of paired bases; asking for structural robustness translates into favouring a high fraction of paired bases; specific motif sequences affect particular interactions with other molecules; certain structural motifs are responsible for ribozyme-like functions, such as the cleaving activity of hammerheads. Properties such as high structural robustness, here understood as the effect of point mutations in the secondary structure [9], might emerge if selection favours the formation of long helices in the structure. Our results lead us to hypothesize that viroid-like replicons might emerge *de novo* with relative ease through a parsimonious process triggered by small RNAs of indistinct origin. The requirement for the process to start is the appearance of a minimal combination of sequence and structural motifs for those RNAs to be recognized by the replication machinery of cells. Sequence elongation and acquisition of new functions might proceed step-wise without major difficulties.

## 2. Materials and Methods

### 2.1. Structural Properties of Extant Viroids

All the viroid species included in this study, as well their accepted taxonomic classification (ICTV 2018b release), are listed in Table 1. For each viroid species, the reference RNA genome sequences were downloaded from NCBI Taxonomy Browser at https://www.ncbi.nlm.nih.gov/Taxonomy/Browser/wwwtax.cgi?id=12884 (last accessed 03/25/19).

**Table 1.**
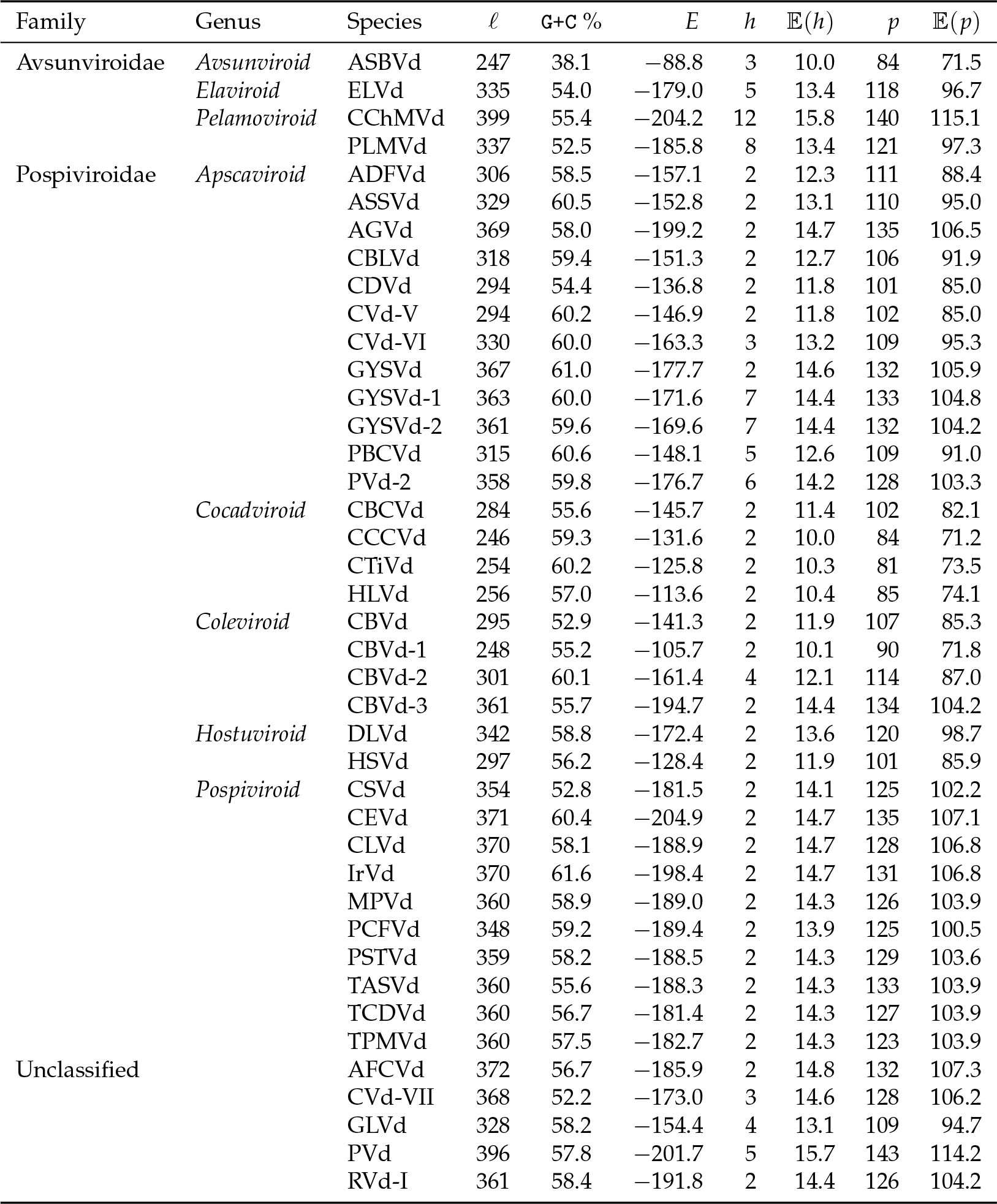
Structural properties of viroids of length *l*. We list their G+C content, the minimal folding energy *E* in kcal mol^*−*1^ (see Materials and Methods), and the number *h* of hairpins and *p* of base pairs. For comparison, the expected values of the number of hairpins, 𝔼(*h*), and base pairs, 𝔼(*p*), in exact calculations of structures of each length (c.f. equation (6)) are reported along the actual values.

We folded the viroid circular RNA sequences into their minimum free energy (MFE) structure using the circfold routine from the ViennaRNA package version 2.1.7 [23], and setting the temperature at 25°C. We then computed the number of hairpins and base pairs, the free energy of the structure, and the G+C content of the sequence (see Table 1).

### 2.2. Computational model

We performed three sets of computational experiments. In the first one, we evolved populations of *N* = 1, 000 circular RNA sequences of fixed length *l* = 300 nt using Wright-Fisher dynamics, letting them evolve for *T* = 20, 000 generations. At each generation, sequences were folded into their MFE as described above. In order to select for fitter sequences, we used the fitness function

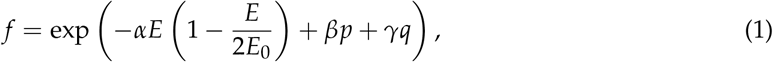

where *E* is the free energy of the folded structure relative to the sequence length *l*, *E*_0_ is a reference energy value, *p* is the number of pairs in the structure and *q* is the number of times a specific sequence motif is found in the sequence. In our simulations, we have taken *E*_0_ = *−*0.433 kcal mol^*−*1^ nt^*−*1^, which is representative of the folding energy per nucleotide of Pospiviroidae (*e.g.* HSVd in Table 1). As for the sequence motif, we have used CNGRRGRRAYCN as an example, which corresponds to the consensus sequence of the periodic motif in CCCVd-small and in PSTVd [20,24]. This fitness function penalizes those sequences whose free energy is far from *E*_0_*l* (when *l* = 300 this is equivalent to penalizing sequences far from *−*130.0 kcal mol^*−*1^). Functions similar to Eq. (1) to select for more than one fitness trait have been used elsewhere [25]. We started all simulations with a randomly sampled seed sequence. Sequences were chosen to reproduce in proportion to their fitness *f*, and mutations were chosen from a Poison distribution of parameter 0.1, which is equivalent to implementing a mutation rate per base pair *µ_P_* = 3.3 *×* 10^*−*3^.

In the second set of experiments, we started the evolutionary dynamics with sequences of length 30 nt. Dynamics were as in the first set of experiments, but an additional mutation mechanism was included: insertions occurred at a rate *µ_I_* = 3.3 *×* 10^*−*3^ per base, like point mutations. In either case, four different situations were analysed: (a) only selecting for energy (*α* = 2.0, *β* = 0.0, *γ* = 0.0); (b) selecting for energy and number of base pairs (*α* = 2.0, *β* = 2.0, *γ* = 0.0); (c) selecting for energy and sequence motifs (*α* = 2.0, *β* = 0.0, *γ* = 20.0); and (d) selecting for energy, pairs and motifs (*α* = 2.0, *β* = 2.0, *γ* = 20.0). Parameters were chosen such that the (one to three) selection pressures acting on the molecules were of similar strength.

In the third set of experiments, we evaluated the likelihood that rod-like molecules of length 130 nt that had evolved under the conditions specified in (d) of the second set of experiments would maintain their folded structure under recombination with a hammerhead ribozyme. For simplicity, the sequence 5’-GAAGAGUCUGUGCUAAGCACACUGACGAGUCUGUGAAAUGAGACGAAACUCUUU-3’, corresponding to the hammerhead ribozyme of avsunviroids [26] was added to different loops (terminal hairpin loops, internal loops or bulges) of the rod-like structures. The complete sequence was subsequently folded into its minimum free energy structure to evaluate if both the paired structure of the evolved rod-like molecule and the hammerhead structure were maintained upon addition of the two modules.

## 3. Results

In order to quantify the ease with which viroid-like properties could emerge in evolutionary scenarios, we have addressed different aspects. First, we have calculated expected features of typical structures for short RNA sequences (of length up to 30 nt) and for sequences with the lengths of the viroids analysed. Second, we have performed numerical simulations of circular RNA populations under a variety of assumptions and compared several properties of evolving populations with those of extant viroids.

### 3.1. Quantitative properties of viroid-like RNA sequences and folds

#### 3.1.1. Structural Properties of Circular RNAs

The folding constraint of circular RNA sequences poses severe biases on the feasibility and abundance of the different structures. Counting how many structures a circular sequence of length *l* can fold into, having *h* hairpins and *p* base pairs, is a nontrivial combinatorial problem that has nevertheless been solved recently with the help of generating functions [27]. If *v*_*l,p,h*_ denotes this number, then

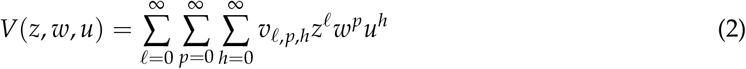

defines its generating function, which turns out to be

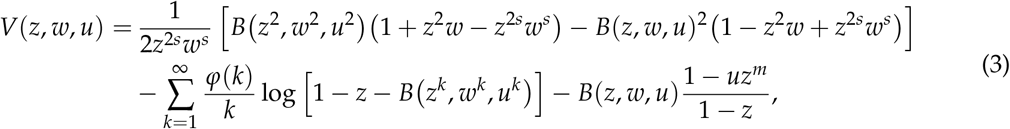

where *ϕ*(*k*) is Euler’s totient function, *B*(*z*, *w*, *u*) solves the equation

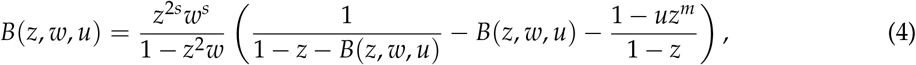

and *m* and *s* are the minimum number of base pairs in a stem and of unpaired bases in a hairpin loop, respectively —two energetic constraints on foldings. (See Ref. [27] for full details.)

Complicated as this may look, it is not difficult to extract asymptotic expressions for *v*_*l,p,h*_ when the sequences are long [27]. The total number of structures grows as

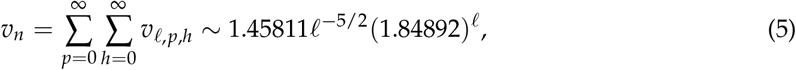

a result analogous to that obtained for the number of structures for open RNA chains [28]. On the other hand, the probability *v*_*l,p,h*_/*v*_*r*_ that a circular structure of length *l* has *p* base pairs and *h* hairpins follows a bivariate normal distribution. In particular, the expected number of base pairs and hairpins grow with *l* as

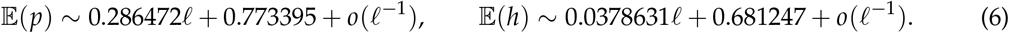

Table 1 lists these expected values for the lengths of extant viroids.

We can further use Eq. (3) to extract the exact count for short sequences using some symbolic algebra package. The results are listed in Table 2 for sequences shorter than 30 bases. It is worth stressing the dominance of rod-like structures for these short sequences. In particular, a third hairpin does not appear until length 21 nt, and even at length 29 nt, the abundance of rods over other structures is nearly seven-fold.

**Table 2.** Number of circular RNA structures of length *l*, exhibiting *h* hairpins and *p* base pairs.

| $\ell$ | $h$ | $p = 2$ | 3 | 4 | 5 | 6 | 7 | 8 | 9 | 10 | 11 |
| --- | --- | --- | --- | --- | --- | --- | --- | --- | --- | --- | --- |
| 10 | 2 | 1 |  |  |  |  |  |  |  |  |  |
| 11 | 2 | 1 |  |  |  |  |  |  |  |  |  |
| 12 | 2 | 2 | 1 |  |  |  |  |  |  |  |  |
| 13 | 2 | 2 | 1 |  |  |  |  |  |  |  |  |
| 14 | 2 | 3 | 2 | 1 |  |  |  |  |  |  |  |
| 15 | 2 | 3 | 2 | 2 |  |  |  |  |  |  |  |
| 16 | 2 | 4 | 3 | 6 | 1 |  |  |  |  |  |  |
| 17 | 2 | 4 | 3 | 10 | 3 |  |  |  |  |  |  |
| 18 | 2 | 5 | 4 | 19 | 9 | 1 |  |  |  |  |  |
| 19 | 2 | 5 | 4 | 28 | 18 | 4 |  |  |  |  |  |
| 20 | 2 | 6 | 5 | 44 | 33 | 16 | 1 |  |  |  |  |
| 21 | 2 | 6 | 5 | 60 | 53 | 36 | 5 |  |  |  |  |
|  | 3 |  |  |  |  | 1 |  |  |  |  |  |
| 22 | 2 | 7 | 6 | 85 | 81 | 82 | 23 | 1 |  |  |  |
|  | 3 |  |  |  |  | 2 |  |  |  |  |  |
| 23 | 2 | 7 | 6 | 110 | 116 | 151 | 64 | 6 |  |  |  |
|  | 3 |  |  |  |  | 7 | 1 |  |  |  |  |
| 24 | 2 | 8 | 7 | 146 | 161 | 276 | 157 | 34 | 1 |  |  |
|  | 3 |  |  |  |  | 20 | 6 |  |  |  |  |
| 25 | 2 | 8 | 7 | 182 | 215 | 452 | 322 | 106 | 7 |  |  |
|  | 3 |  |  |  |  | 42 | 21 | 2 |  |  |  |
| 26 | 2 | 9 | 8 | 231 | 281 | 731 | 614 | 294 | 45 | 1 |  |
|  | 3 |  |  |  |  | 84 | 56 | 14 |  |  |  |
| 27 | 2 | 9 | 8 | 280 | 358 | 1106 | 1068 | 665 | 166 | 8 |  |
|  | 3 |  |  |  |  | 156 | 126 | 57 | 4 |  |  |
| 28 | 2 | 10 | 9 | 344 | 449 | 1652 | 1773 | 1401 | 507 | 60 | 1 |
|  | 3 |  |  |  |  | 264 | 252 | 176 | 28 |  |  |
|  | 4 |  |  |  |  |  |  | 1 |  |  |  |
| 29 | 2 | 10 | 9 | 408 | 553 | 2360 | 2783 | 2668 | 1279 | 248 | 9 |
|  | 3 |  |  |  |  | 429 | 462 | 456 | 130 | 5 |  |
|  | 4 |  |  |  |  |  |  | 2 |  |  |  |

#### 3.1.2. Phenotype sizes

The abundance of sequences yielding structures with specific properties cannot be directly estimated from Table 2, since the size of a phenotype (meaning in this case the total number of sequences folding into a specific shape) depends on its structural properties [29]. For example, the higher the number of pairs in an RNA secondary structure, the fewer the number of sequences compatible with such a structure [30]. Furthermore, structures with a low number of pairs are energetically disfavoured [31], since stabilizing a structure with short stacks requires pairs of low free energy such as `G-C`, and this condition limits the number of sequences compatible with such structures. On the other hand, paired nucleotides have lower neutrality than unpaired ones (they admit fewer changes without modifying their paired condition) [32,33], so structures with many paired positions tend to have smaller phenotypic sizes. Eventually, it turns out that typical structures (the most abundant ones in Table 2) also have the largest phenotypes, so that the frequency of sequences folding into typical structures is much higher that the frequency of typical structures themselves. While the latter quantity can be derived from Table 2, the former requires full consideration of the sequence-to-MFE secondary structure map.

Our analytical results indicate that viroid-like folds, and especially rod-like shapes (those with *h* = 2), become increasingly rare in the space of structures as sequence length grows. Equation (6) shows that the number of hairpins increases approximately one unit every 26 nt in the sequence. Still, the huge degeneracy between RNA sequence and structure predicts an astronomically large number of sequences folding into a vast majority of structures (including those of viroids). Results in [34] based on the computational exploration of the sequence-to-structure map in RNA [35] allow us to estimate the size *S* of any RNA structure with 2*p* paired nucleotides and *u* = *l −* 2*p* unpaired nucleotides as

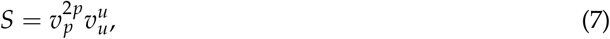

where *v*_*p*_ = 1.17 ± 0.08 and *v*_*u*_ = 2.79 0.08 are numerically obtained quantities [34]. The number of sequences compatible with viroid structures varies between about 10^46^ for CCCVd and 10^72^ for CChMVd. However, note that a typical structure of length 246 nt harbors about 142 paired nucleotides, so its phenotype size becomes of order 10^56^. That size rises to about 10^91^ for a typical structure of length 399 nt (see Table 1 for a comparison between the number of pairs in each viroid and the number expected in random sequences of the same length).

#### 3.1.3. Probability of specific sequence motifs in random sequences

It is well known that specific sequences in the viroid genome are related to functions that are essential to complete the viroid cycle. Such sequences have been identified in PSTVd to promote entry into the nucleus [36], in HSVd related to pathogenicity [37], in the central conserved region (CCR) of different *Pospiviroid* to guarantee optimal replication [38] or in the hammerhead motif of Avsunviroidae for effective autocatalytic activity [39]. Indeed, viroid sequences are subjected to a diversity of selective pressures that lead to conserved positions and regions [7].

In general, the likelihood of appearance of a specific sequence motif can be estimated as follows. As an illustration, consider CNGRRGRRAYCN, the consensus sequence of the periodic motif in CCCVd-small and in PSTVd [20,24], which will be later used in the numerical simulations. The calculation proceeds in the same fashion for any other case. This sequence has 5 fixed positions, 2 positions that can be occupied by any nucleotide, 4 positions by a purine, and 1 by a pyrimidine. There are *N* = 4^2^ × 2^4^ × 2^1^ = 2^9^ possible sequences out of 4^12^ = 2^24^ different ones. The likelihood that the motif CNGRRGRRAYCN appears at a fixed position in a sequence (that is, assuming that there is a unique possible site for the initial C) is 2^*−*15^ *≈* 3 *×* 10^*−*5^; in other words, the motif appears on average in 3 out of 100, 000 molecules. However, if the initial position does not matter, this number significantly grows with the length of the molecule. Note that the probability that the motif appears at least once anywhere in a sequence of length *l* is 1 *−* (1 *−* 2^*−*15^)^*l*^ ≈ 2^*−*15^*l*. For random sequences of length *l* = 300, the motif will be found in almost 1% of the sequences.

The random appearance of a circular RNA sequence folding into a rod-like structure with a sequence motif that promotes interaction with other molecules (*e.g.* polymerases, RNases or ligases) is therefore not just a possible event, but a highly likely one. The stochasticity inherent to this random matching between two dissimilar molecules might perhaps explain why Pospiviroidae use DNA-dependent RNA polymerase II instead of a nuclear RNA-directed RNA polymerase [2], or why PSTVd uses DNA ligase 1 to circularize the genomic RNA monomers [40]. Actually, the results above suggest that, in view of the ubiquity of circular RNAs, potential viroid-like replicons are steadily generated, in the absence of any specific selection pressure, with high likelihood.

### 3.2. Evolutionary Dynamics of Circular RNAs of Fixed Length

*In silico* evolutionary experiment with populations of circular RNAs of size 300 nt have been performed with the aim of addressing two main questions: Which secondary structures dominate in such populations when different selection pressures are applied? How likely is it to evolve from *Pospiviroid*-like structures to *Avsunviroid*-like structures (or the other way round)?

In all numerical experiments performed, an average energy of the folded structure similar to that observed in natural viroids (of similar length) was preferentially selected. This selection pressure responds to the observation of viroids maintaining a sufficiently low energy so as to fold into stable structures but sufficiently high so as to be opened with relative ease for replication. A second selection pressure applied aims at maximizing the number of pairs with the goal of favouring structural robustness. If this pressure is applied in the absence of selection for an average energy, the G+C content increases without restrictions, leading to highly stable and robust structures that, however, do not show any plasticity and, as a consequence, would be very difficult to replicate. A third selection pressure regards a specific sequence motif, where we have chosen CNGRRGRRAYCN as an example. For simplicity, let us represent by *P* (*M*) the situation where increasing the number of pairs (minimizing the distance to the sequence motif) increases fitness, while selection for an average energy *E* occurs in all cases.

Figure 1 summarizes the results of the four situations once the populations have reached a statistically stable equilibrium. Colored curves correspond to numerical simulations (*E*, blue curve; *E* + *M*, orange curve; *E* + *P*, green curve; *E* + *P* + *M*, red curve), while grey histograms correspond to extant viroids and are displayed for comparison. The four situations can be grouped into two major qualitative behaviours: *E* resembles *E* + *M* and *E* + *P* is similar to *E* + *P* + *M*.

**Figure 1.**
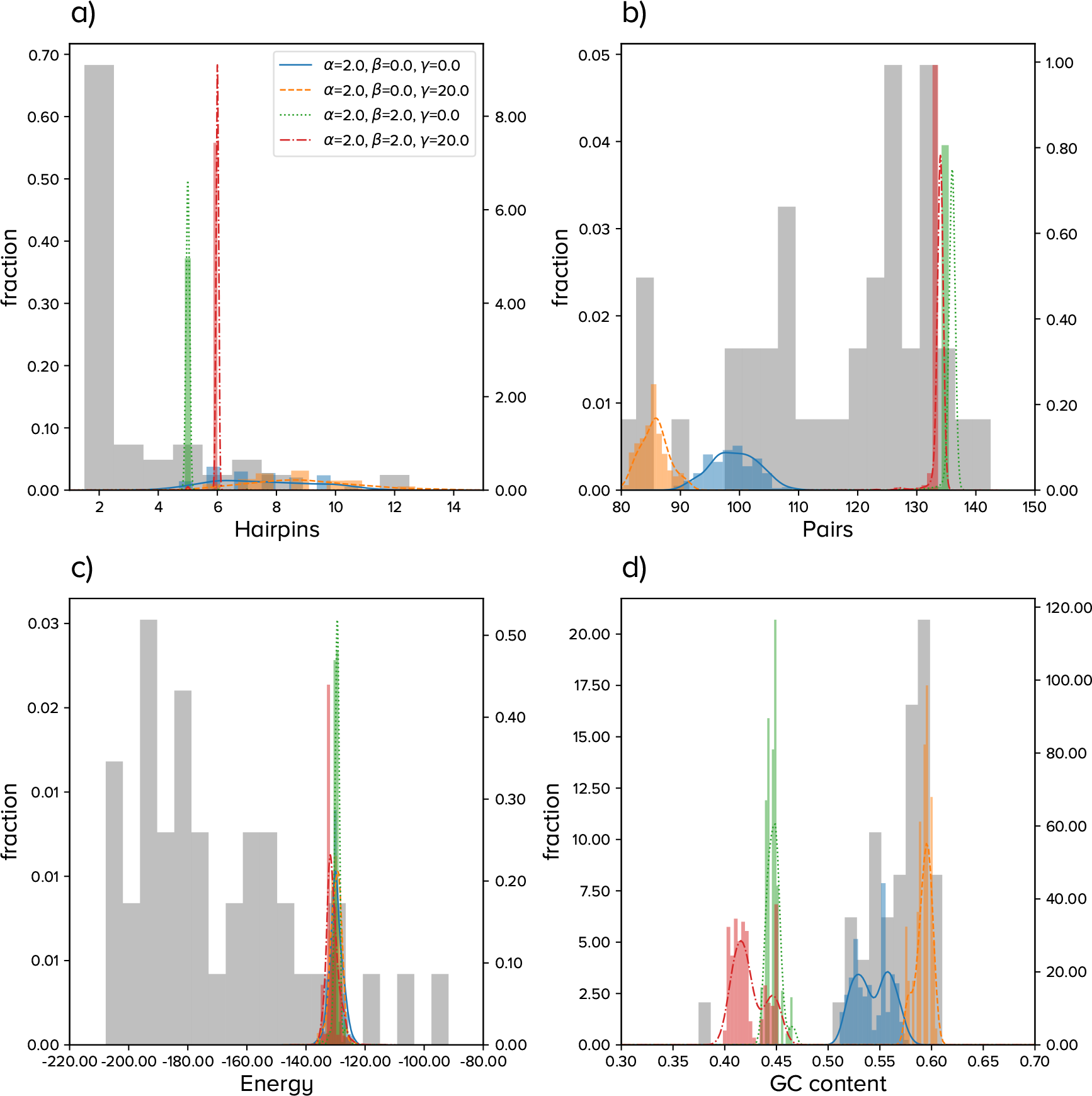
Summary of results for the evolution of populations of circular RNA sequences of fixed length. Histograms in color correspond to numerical simulations with parameters specified in the legend. Lines are kernel density estimates of the underlying distribution. Grey bars correspond to viroids in Table 1. (**a**). Number of hairpins *h*; (**b**). Number of base pairs *p*; (**c**). Minimum free energy of the secondary structure; (**d**). `G+C` content.

If only energy or energy and a sequence motif are positively selected, we obtain populations highly heterogeneous in structure, with a broad distribution for the number of hairpins (from 4 to 12, with typical numbers around 6-9). Also, the number of paired nucleotides follows broad distributions, with averages that keep relatively low as compared to most viroids. In order to maintain the average energy required in the simulations, and as a result of the latter, the G+C content attains high values.

The high diversity of the former populations severely decreases when selection favouring the increase in the number of pairs is considered. This selective pressure leads to populations with a lower number of hairpins (with average around 5 or 6), a significantly larger number of pairs and, in agreement, a lower G+C content. Note that a sequence of length *l* with *h* hairpins (whose minimum size is 3) has a maximum number of pairs *p*_mx_ = (*l −* 3*h*)/2. For *l* = 300 and *h* = 5, *p*_mx_ = 142, so the number of pairs is close to maximal in these simulations. Our results show a relationship among number of pairs, folding energy, and G+C content. If *E*_0_ decreases, and since the number of pairs is already close to its highest possible values, the G+C content has to increase to meet the requirement. This would lead to sequences more similar to actual viroids, but even less plastic that those found in our simulations and holding a very low structural diversity in their populations. If *E*_0_ decreases too much, the opening of those structures to perform the necessary functions via interaction with cellular components would be compromised.

Finally, we have monitored the number of appearances of the sequence motif CNGRRGRRAYCN in all four situations (results not shown). When the motif is selected for, it appears almost immediately in the situation *E* + *M*, while it takes a few thousand generations to emerge under *E* + *P* + *M*. The number of repetitions per sequence grows and reaches over 20 appearances per sequence in *E* + *M*, and about 3 appearances per sequence in *E* + *P* + *M* by the end of our simulations, though this number does not seem to have saturated. These results agree with the huge degeneracy of the sequence-to-structure map, which permits, to a good extent, the simultaneous and successful selection for structure and specific sequence motifs.

The numerical results reported in this section suggest that selection of rod-like shapes is unlikely if evolution starts with relatively long sequences. Requiring a high number of pairs without limiting the number of hairpins yields long helices but branched structures, with no less than 5 hairpins and a low population diversity. An example of the evolutionary dynamics in the situation *E* + *P* + *M* is represented in Figure 2 (a) and (c). Moreover, if the number of base pairs is positively selected, and independent of other selective pressures, long stacks are locally fixed, since a large number of paired bases is however compatible with highly branched structures. The effect of point mutations on structures with long helices is typically small, such that populations are trapped in structure space and minor changes in sequence find it hard to modify the overall structure [9,41], hence the number of hairpins. Our results support previous suggestions that viroids might be evolutionary trapped due to adaptive constraints [8]. Here we show that maximizing the number of pairs is indeed a constraint limiting innovation if only minor changes in sequence occur. As a side result, it seems implausible that the two viroid families are phylogenetically linked at a late evolutionary stage, that is, for sequence lengths comparable to those of extant viroids. In the same vein, the emergence of viroid-like parasites through evolution of the large circular RNAs expressed in eukaryotes might be difficult [42,43].

**Figure 2.**
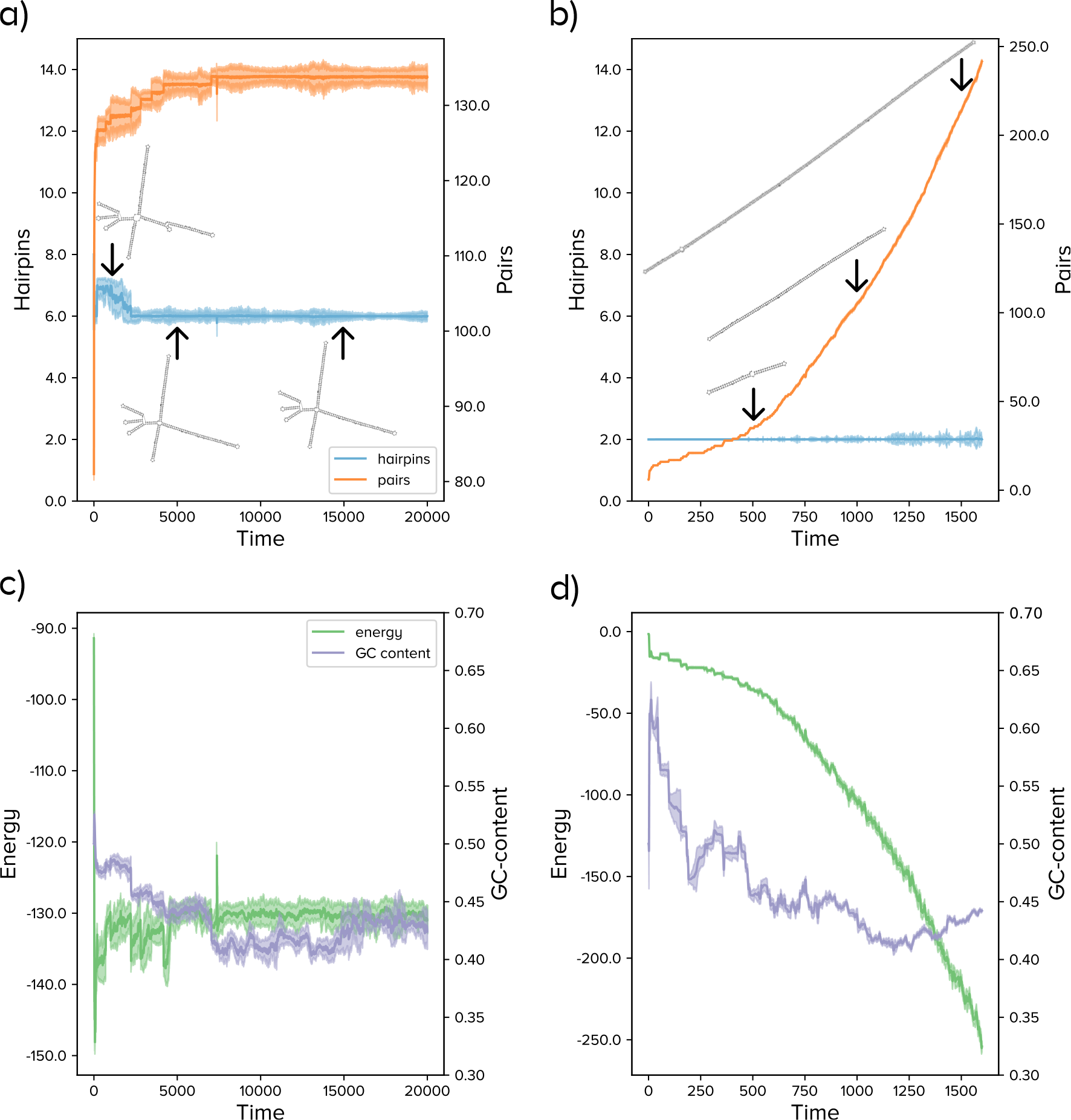
Evolutionary dynamics of RNA populations with fixed (a,c) and variable (b,d) length. In the former case, simulations start with random sequences of length 300 nt; in the latter, of length 30 nt. (**a**). The number of hairpins decreases initially but rapidly freezes around 6; the number of base pairs increases until a value near the maximum possible for that number of hairpins. The most abundant structure is shown at three different time point, as indicated by arrows; (**b**). As in (a), for sequences growing in length; (**c**). Dynamics of energy and G+C content for the same run shown in (a); (**d**) As in (c), for the run shown in (b).

### 3.3. Evolutionary Dynamics of Circular RNAs of Increasing Length

The possibility that extant viroids have evolved from large circular RNAs, either random or selected for other functions, seems unlikely. Trapping in configuration space might play a two-fold important role: evolution of an RNA molecule by point mutations might involve too large evolutionary times, while at the same time it serves to preserve the original function. In particular, and as we have just shown, the structural plasticity of viroid-like structures, with many base pairs, is lower than average (compared to random sequences of the same length). Perhaps a more plausible evolutionary pathway for the emergence of viroid-like replicons would start with random short sequences whose length might increase concomitantly with functional selection. Among others, micro-RNAs (miRNA) [44] could be potential candidates to trigger the process that we describe in this section. Intriguingly, miRNAs and viroids have been found to share important structural features [45], among which stands the pervasive presence of stem-or rod-like secondary structures, respectively.

Numerical and analytical calculations demonstrate that hairpins [27,31,46] and small rods ([27] and this work) are highly preferred structures for short RNAs, so that these shapes are dominant in the absence of specific selection pressures. Circularization of short ssRNAs should occur frequently, among others as a spontaneous product of the ligase activity exhibited by hairpin structures [47,48]. Notably, a ribozyme activity of hairpin-like structures was first described in plant virus satellite RNAs [49,50].

We start the simulations in this section by taking as initial condition a population of circular RNAs of length 30 nt. They overwhelmingly fold into rod-like structures. Selection pressures applied are as in the former section. We analyse two evolutionary mechanisms: addition of nucleotides and recombination with an independent hammerhead ribozyme. The addition of simple sequence repeats is an additional plausible mechanism for elongation, consistent with observations [51], that we do not consider explicitly here. The main question addressed is whether the rod-like structure that seeds the process is maintained along evolution.

#### 3.3.1. Growth Through Insertion of Single Nucleotides

Simulations for growing sequences have been repeated in the four situations described, *E*, *E* + *M*, *E* + *P* and *E* + *P* + *M*. As above, qualitative results group into two different pairs: *E* is akin to *E* + *M*, while more interesting results are obtained for *E* + *P* or *E* + *P* + *M*. If selection for the number of pairs is absent, sequences elongate at a slow pace initially, apparently more as a result of drift than as a consequence of any of the selective pressures acting. After evolution for several tenths of generations, sequence lengths reach sizes that vary between 300 nt and 500 nt, a growth that is accompanied by a significant increase in the number of hairpins. The number of base pairs is well below its maximum possible value, meaning that loops are frequent and/or large: populations are mainly formed by highly unstructured molecules of low structural robustness. As expected, sequence motifs are absent in the *E* situation (they occasionally appear but are not fixed) and reach 15 repetitions (and growing) after about 6 *×* 10^5^ generations in the *E* + *M* situation.

The results of simulations for the situation *E* + *P* + *M* (qualitatively equivalent to *E* + *P*) are summarized in Figure 2 (b,d), and compared with our numerical results in populations of sequences of constant length. At odds with what we observed in the latter case, the gradual addition of nucleotides preserves the initial number of hairpins, and therefore yields increasingly large rods. Examples of the most abundant structures at different time points are shown in Figure 2 (b), as indicated by arrows. The `G+C` content of the sequences stabilizes at around 0.4-0.45 (compare with the red curve in Fig 1 (d)), while the minimum folding energy decreases proportionally to the number of pairs. In just 10^3^ generations, the populations are ensembles of rod-like structures with size comparable to that of extant viroids and at least one copy of the sequence motif under selection.

#### 3.3.2. Growth through Modular Recombination

Some authors have described viroid structure as a “collection of structural motifs which play specific functional roles in viroid replication, processing, transport, and pathogenesis” [7]. Though not explicitly discussed, this modular structure hints at the possibility that the different functional abilities of viroids could have been acquired through recombination with functional RNAs of different origins. Viroids formed through recombination of fragments present in other viroids have been described [52, 53]. A highly plausible case of modular recombination is provided by Hepatitis *δ* virus (HDV) [54], the unique animal pathogen described to date with a viroid-like non-coding domain [55] and a second domain coding for an antigen of independent evolutionary origin [56]. The possibility that independent RNA modules, functional in other molecular contexts, could have endowed *bona fide* viroids with new functions remains however as a hypothesis. Still, modular evolution has several advantages (as compared to direct evolution of longer molecules) [57] which may have been determinant in the early emergence of chemical function [46].

As a representative example, we quantify here the likelihood that important functions of viroids dependent on their secondary structure would be preserved under modular evolution. First, we have evolved single sequences as in the previous section until they have reached length 130 nt. In all cases, they had a rod-like secondary structure, but there were variations, albeit narrow, in their energy, `G+C` content or number of base pairs. At that point, we studied the effect of ligation of the evolved sequence to the hammerhead ribozyme of viroids [26] (see Materials and Methods). Figure 3 portrays the structure of the hammerhead and summarizes the possible outcomes of the process. The hammerhead structure is preserved in 12% of recombination events (two possible cases are illustrated in Fig. 3 (c) and (d)). Still, the two structures are relatively easy to preserve, for instance if recombination occurs at one of the terminal hairpins. Specifically, we found that this happens in 8% of cases, Fig. 3 (c).

**Figure 3.**
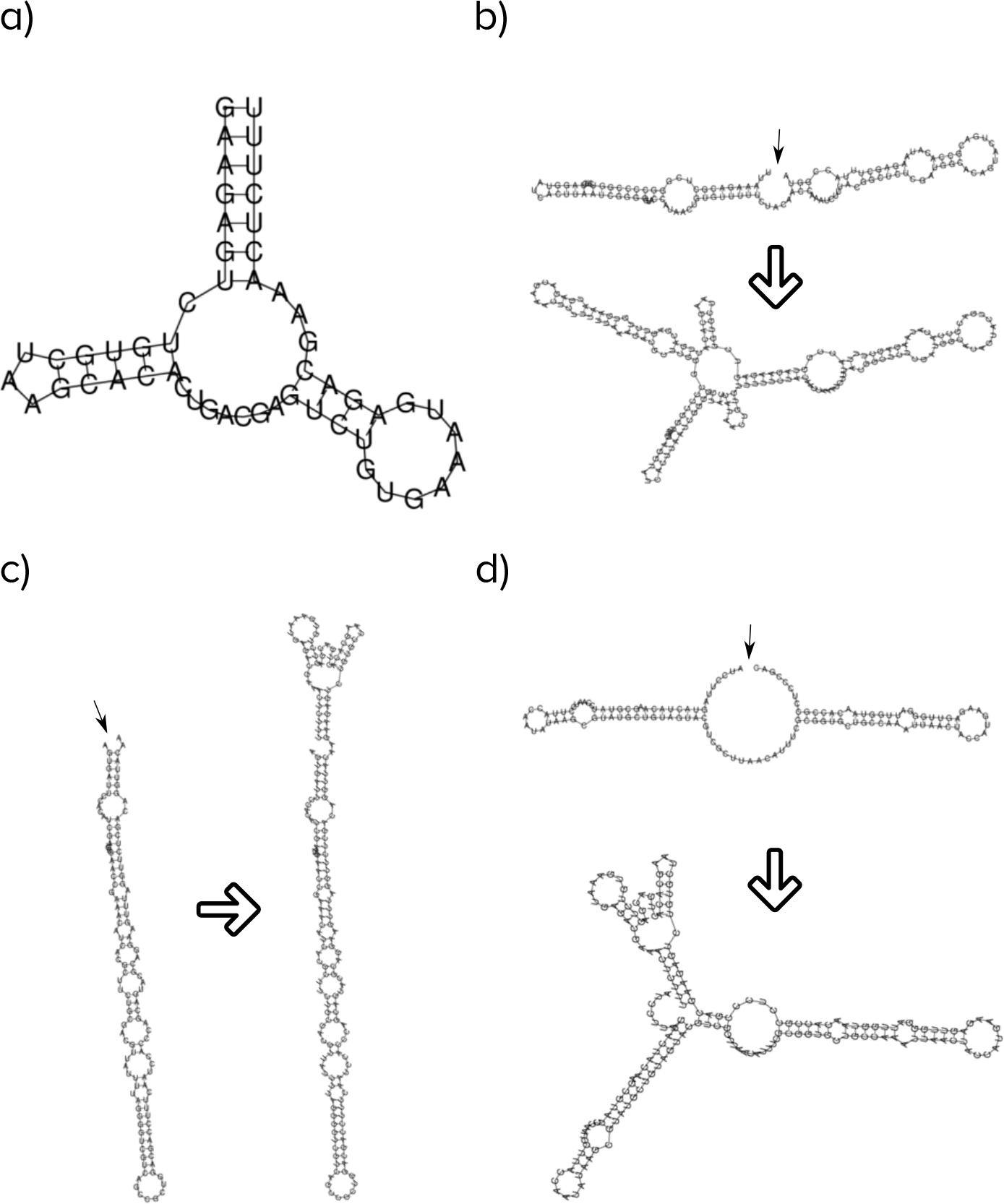
Effects of modular evolution in the structure of rod-like folds. Small arrows signal the recombination site; large arrows indicate the transformation of the structure upon recombination with the hammerhead. (**a**). Structure of the example hammerhead to be recombined with evolved sequences; (**b**). Part of the structure is maintained, part disrupted. The hammerhead structure is lost; (**c**). Both recombining structures are preserved; (**d**). The hammerhead structure is preserved, but the rod-like fold is partly disrupted.

It is likely that the very rod-like structure plays a role towards increasing its preservation under recombination, in the same sense that such structures are more robust to mutations [9,41]. In a different context, the modular combination of two RNA structures resulted in 4% of ligation event preserving the structures of the independent modules, which had a lower fraction of paired bases [57].

## 4. Discussion

The quantitative results reported in this work give support to a *de novo* emergence of viroid-like replicons. We have envisaged a parsimonious scenario, summarized in Figure 4, where short RNA molecules of various origins could circularize and act as seeds of the process. There is no particular requirement in the initial condition: plant and animal cells, in particular, hold a large and variable pool of RNAs fulfilling different functions, of a variety of lengths and origins, which might in practice serve as a test bed for new functions: “A truly modern RNA World exists in each cell” [22]. Circular RNAs are ubiquitous in nature [58] and pervasively expressed in higher eukaryotes [42]. The likelihood that one such RNA bears a specific sequence that could be mistakenly recognized by an RNA polymerase is high. This recognition would be further facilitated by a compact folding mimicking dsDNA [4]. Even the template-free synthesis of RNA can be possible in certain environments [59]. In a prebiotic scenario, random RNA sequences could have fulfilled these minimal conditions, such that viroid-like replicons could have easily emerged in a precellular context. But there are also several different extant RNAs which could be involved in this particular molecular mimicry, a prominent example being miRNAs. The compact, hairpin-like structures of miRNAs, and their high diversity [44] makes of them good candidates to trigger such a process. Regardless of its origin, an RNA molecule that can be replicated in that way would become more abundant in front of other variants, starting in this way its Darwinian evolution towards becoming a fully fledged selfish replicator. This nonetheless, the eventual success of such incipient replicator can be compromised if other abilities are not developed timely. First, its replication would be initially limited to the cellular environment: acquiring the ability to move to neighbouring cells seems a necessary requirement. Second, *de novo* replicators have to persist in a molecular ecology that might prevent its fixation in a variety of ways. In particular, if the potential niche of viroid-like replicons is already occupied, it might be extremely difficult to invade the system and displace the established molecule. The notion that niche occupation limits the success of invasions attempted by ecologically analogous species is widespread in ecology [60,61] and should be applicable, *mutatis mutandis*, to molecular ecologies.

**Figure 4.**
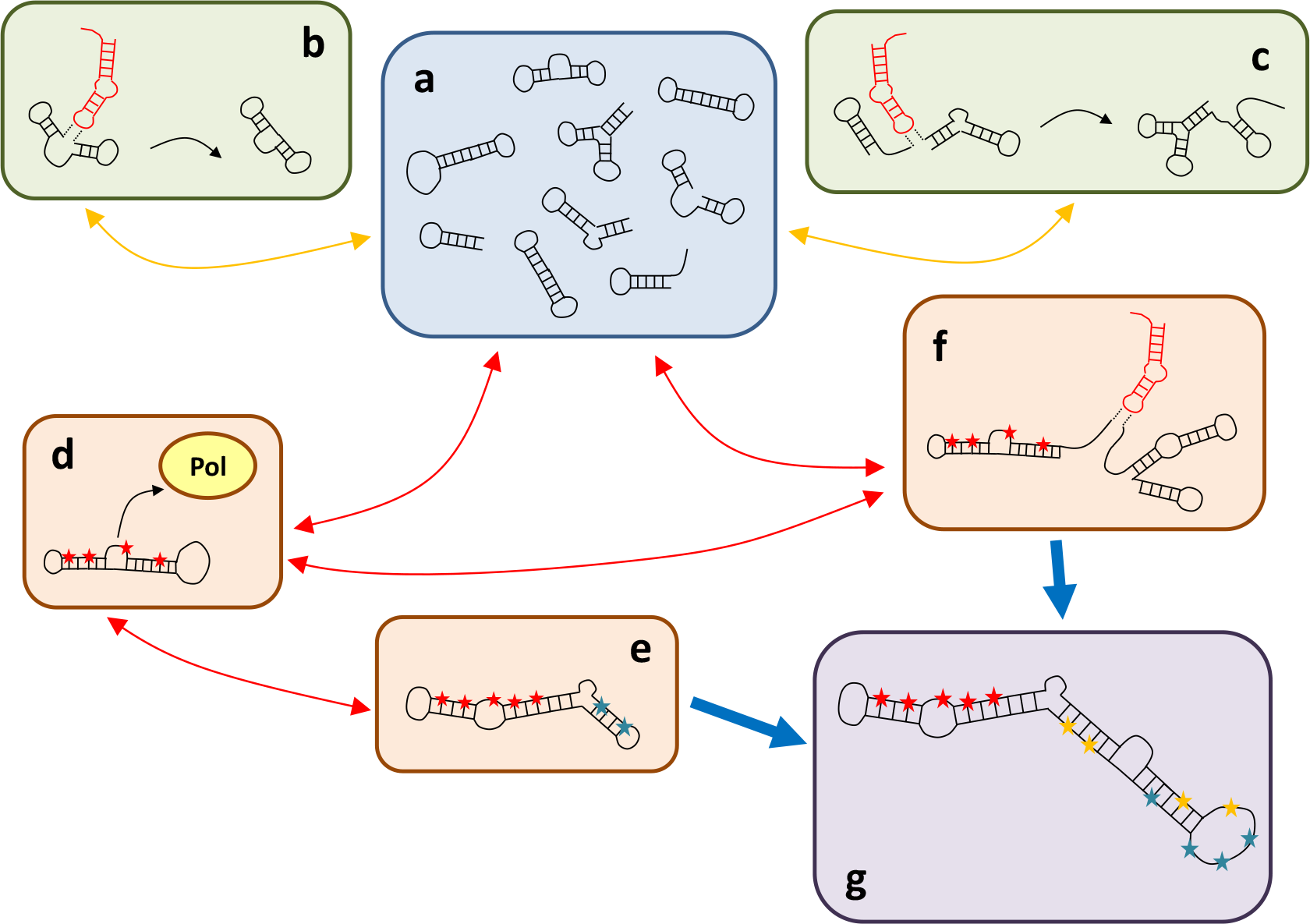
Schematics of a parsimonious scenario leading to the emergence of viroid-like replicons *de novo*. Structures in black represent circulating RNAs with various structures and functions; structures in red stand for hairpins with RNA ligase activity; stars indicate positions in the sequence which are fixed, different colors corresponding to possible motifs that interact with different molecules. (**a**). Circulating pool. There is a pool of RNA sequences of various origins. Short random sequences spontaneously fold into hairpin structures; (**b**). Circularization. Hairpins are able to catalyze ligation reactions, so a fraction of open chains would close in presence of hairpins; (**c**). Modular growth. Independent RNAs can ligate through reactions analogous to those causing circularization. Both reactions in (**b**) and (**c**) yield novel molecules that add to the circulating pool; (**d**). A non-negligible fraction of circulating RNAs might have specific sequence positions that promote interactions with a polymerase. Once this process starts, selection for improved replication is triggered. Those RNAs would become more prevalent in the circulating pool; (**e**). Minimal circular replicons might grow in length through the random addition of nucleotides to their sequences. Sequence motifs with no specific function can evolve to improve the replicative ability of the molecule (*e.g.* by increasing mobility or selecting additional positions to interact with the polymerase); (**f**). New functions can be acquired through recombination of functional RNAs in the circulating pool; (**g**). Sufficiently long replicons that may arise from processes as those in (**e**) and/or (**f**) might respond to a variety of selection pressures. In practice, these replicons occupy a niche in the molecular ecology equivalent to that of viroids.

In any way, subsequent evolution of short replicating RNAs could have occurred through several different, not mutually excluding mechanisms. One is elongation through the addition of stretches of nucleotides. In Spiegelman’s evolutionary experiment, where the unique selective pressure applied favoured faster replication, the length of RNA replicators decreased through evolution [21]. However, this reduction does not need to occur in more complex environments, where other selective pressures might be acting. In some of our *in silico* evolution experiments, we have monitored the appearance of multiple sequence motifs. In a natural environment, it cannot be discarded that initiating replication at more than one site confers an advantage that compensates for the mutational cost of replicating longer molecules. Also, longer sequences can respond to a high number of selection pressures, thus paving the way for the emergence of specific sequence motifs able to fulfill new functions: interaction with other molecules, cell-to-cell movement or improved replication through additional structural motifs are a few examples. Similarly, the duplication of parts of the sequence through imperfect rolling circle replication can be sources of novelty through neo- or subfunctionalization, as it happens with gene duplication. Examples of such processes of genome length increase have been previously documented in the coleviroids [62], the genesis of CCCVd variants containing duplicated segments of the left terminal domain and part of the adjacent variable domain [63] and the long CEVd isolate D-104 [64].

Modular evolution appears as a plausible and highly efficient mechanism to acquire new functions. Repeated recombination between functional modules selected in different contexts [57] could have facilitated the transmission of ubiquitous functions such as cleavage through hammerheads, ligation catalysis through hairpins or long-distance movement motifs. In this context, one wonders whether the similarity in sequence and structure of different viroids and viroid-related replicons has to be interpreted as a result of descent with modification or of horizontal transfer. In the latter case, functional submolecular elements could be better described as the nodes of a network that underlies and favors the rapid emergence of new viroid-like replicons through shuffling of functional modules. This idea has sound support in viruses [65] and may underlie the evolutionary emergence of multipartite viral genomes [66]. Examples of such recombinant origin in viroids are best illustrated by CLVd [52], which resulted from the intracellular recombination between a *Hostuviroid* and a *Pospiviroid* coinfecting the same plant, and AGVd, which results from recombination of GYSVd-1 (an *Apscaviroid*) and CEVd (a *Pospiviroid*) both infecting grapevine plants [53].

The emergence of genetic parasites is unavoidable [67,68]. In the RNA World, primitive selfish replicons must have emerged, and they might well have been viroid-like. However, the ease with which this kind of parasites seems to arise suggests that this replicative strategy might have been discovered multiple times in evolution. Retroviroid-like element provide indirect support to this hypothesis, since they should have appeared following the evolutionary discovery of DNA [4]. Though it has been commonly accepted that Pospiviroidae and Avsunviroidae had a common phylogenetical origin after [55], the cell nucleus and chloroplasts offer significantly different environments wherein viroid-like parasites could proliferate [69]: an independent origin of the two families cannot be discarded in the context here discussed. In an RNA World, chemistry had a completely different nature, DNA was absent and all current proteins did not exist as such. Extant viroids may resemble in many ways early replicons in an RNA World, but this similarity does not imply that the former are linked by descent to the latter (see E. V. Koonin’s comment in [70]). The intimate relationship between viroid sequences and the extant molecules they interact with, together with their fast adaptive responses to new selection pressures speaks for the implausibility of maintaining specific sequences unaltered for billions of years. Though the viroid-like niche is inborn to self-maintained systems, extant viroids are, in all likelihood, *de novo* RNAs.

## Author Contributions

Conceptualization, JAC and SM; methodology, PC, JAC, SM; software, PC and JAC; validation, PC, SFE, JAC and SM; formal analysis, PC and JAC; investigation, PC, SFE, JAC and SM; resources, PC, SFE, JAC, SM; data curation, PC, SFE, JAC and SM; writing—original draft preparation, SM; writing—review and editing, PC, SFE, JAC, SM; visualization, PC and SM; supervision, SM; funding acquisition, PC, SFE, JAC and SM.

## Acknowledgments

The authors want to thank Ronny Lorenz for his assistance on using the ViennaRNA C library.

## Abbreviations

The following abbreviations are used in this manuscript. We list as well the NCBI accession numbers of viroid sequences:

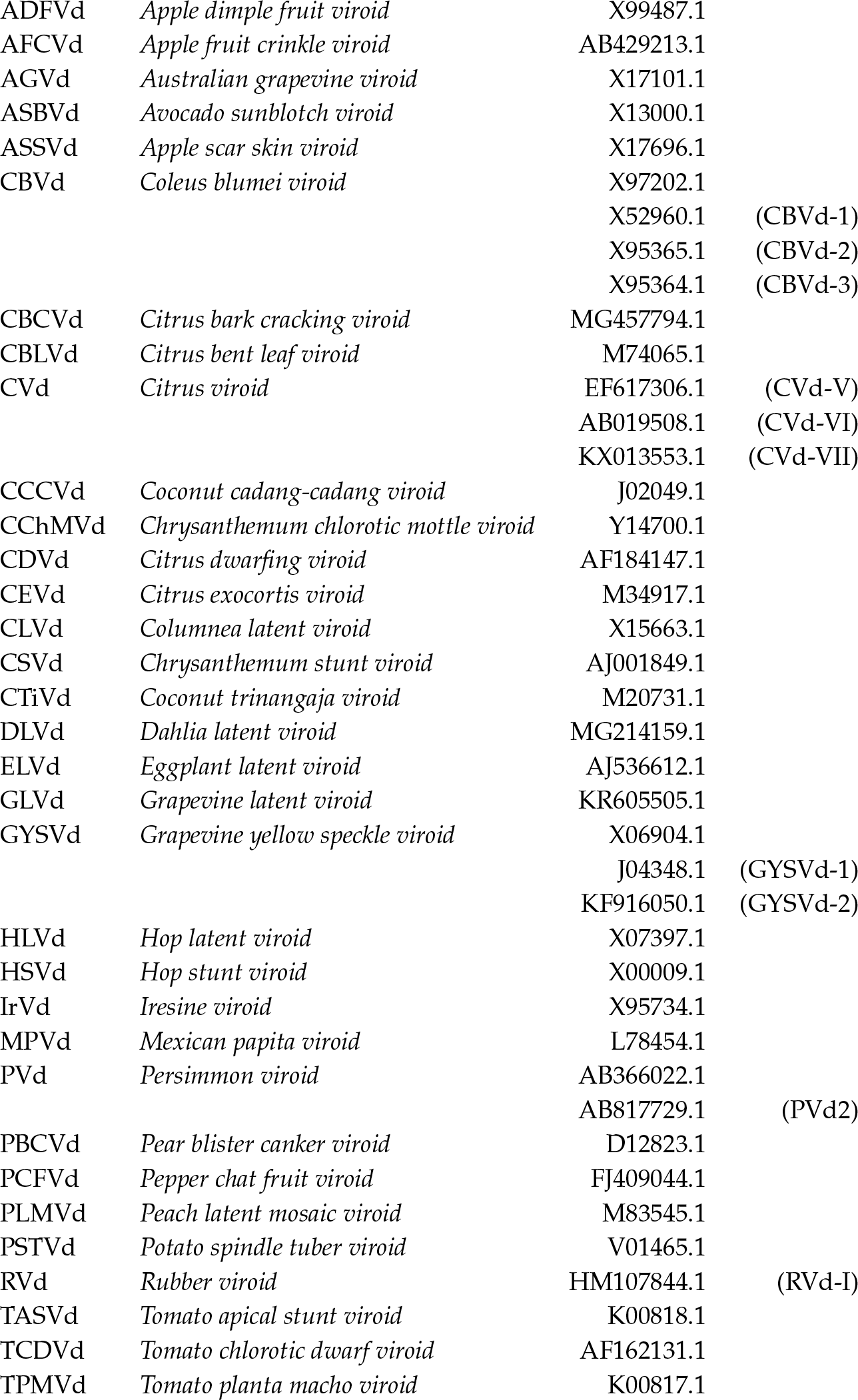

